# A randomised controlled trial of an Intervention to Improve Compliance with the ARRIVE guidelines (IICARus)

**DOI:** 10.1101/370874

**Authors:** Kaitlyn Hair, Malcolm Macleod, Emily Sena, Study steering committee, David Howells, Philip Bath, Study management committee, Cadi Irvine, Catriona MacCallum, Gavin Morrison, Alejandra Clark, Gina Alvino, Michelle Dohm, Programming and data management, Jing Liao, Chris Sena, Redactions, Rosie Moreland, Design of outcome assessment platform, Fala Cramond, Cadi Irvine, Jing Liao, Gillian L. Currie, Zsanett Bahor, Paula Grill, Alexandra Bannach-Brown, Outcome assessment, Kaitlyn Hair, Daniel-Cosmin Marcu, Sarah Antar, Cadi Irvine, Katrina Blazek, Timm Konold, Monica Dingwall, Victoria Hohendorf, Mona Hosh, Paula Grill, Klara Zsofia Gerlei, Kimberley Elaine Wever, Emily Sena, Victor Jones, Terence J Quinn, Natasha A Karp, Jennifer Freymann, Anthony Shek, Teja Gregorc, Arianna Rinaldi, Privjyot Jheeta, Ahmed Nazzal, David Ewart Henshall, Joanne Storey, Julija Baginskaite, Cilene Lino de Oliveira, Kamil Laban, Emmanuel Charbonney, Savannah A. Lynn, Marco Cascella, Emily Wheater, Daniel Baker, Gillian L. Currie, Ryan Cheyne, Edward Christopher, Paolo Roncon, Evandro Araújo De-Souza, Mahmoud Warda, Sarah Corke, Zeinab Ammar, Leigh O’Connor, Ian M. Devonshire, Reconciliation, Kaitlyn Hair, Daniel-Cosmin Marcu, Sarah Antar, Timm Konold, Monica Dingwall, Emily Sena, Paula Grill, Sarah K. McCann, Data analysis, Jing Liao, Laura J Gray, Ezgi Tanriver Ayder, Writing committee

## Abstract

The ARRIVE (Animal Research: Reporting of In Vivo Experiments) guidelines are widely endorsed but compliance is limited. We sought to determine whether journal-requested completion of an ARRIVE checklist improves full compliance with the guidelines. In a randomised controlled trial, manuscripts reporting *in vivo* animal research submitted to PLOS ONE (March-June 2015) were allocated to either requested completion of an ARRIVE checklist or current standard practice. We measured the change in proportion of manuscripts meeting all ARRIVE guideline checklist items between groups. We randomised 1,689 manuscripts, 1,269 were sent for peer review and 762 accepted for publication. The request to complete an ARRIVE checklist had no effect on full compliance with the ARRIVE guidelines. Details of animal husbandry (ARRIVE sub-item 9a) was the only item to show improved reporting, from 52.1% to 74.1% (X^2^=34.0, df=1, p=2.1×10^−7^). These results suggest that other approaches are required to secure greater implementation of the ARRIVE guidelines.

## Background

There are widespread failures across *in vivo* animal research to adequately describe and report research methods, including critical measures to reduce the risk of experimental bias (Kilkenny et al., 2009, Macleod et al., 2015). Such omissions have been shown to be associated with overestimation of effect sizes (Macleod et al., 2015, Hirst et al., 2014) and are likely to contribute, in part, to translational failure. In an effort to improve reporting standards, an expert working group coordinated by the National Centre for the Replacement, Refinement and Reduction of Animals in Research (NC3Rs) developed the Animal Research: Reporting of In Vivo Experiments (ARRIVE) guidelines (Kilkenny et al., 2010), published in 2010.

Since the ARRIVE guidelines were first published, they have been endorsed by many journals in their instructions to authors, but this has not been accompanied by substantial improvements in reporting (Baker et al., 2014, McGrath and Lilley, 2015, Gulin et al., 2015, Avey et al., 2016). Simply endorsing the guidelines does not appear to be sufficient to encourage compliance. Recent findings suggest that following the introduction of mandated completion of a distinct reporting checklist at ten Nature Journals at the stage of first revision significantly improved the quality in reporting versus that of comparator journals (Han et al., 2017, Macleod, 2017)

PLOS ONE is an open access online only journal which at the time this study began published around 32,000 research articles per year. Of these, some 5,000 described *in vivo* research. At present, PLOS ONE instructions to authors encourage compliance with the ARRIVE guidelines, but do not mandate checklist completion. Journals have an important role to play in ensuring that the quality of reporting in the research they publish is robust, yet the most effective mechanism by which they can achieve this remains unclear.

Our aim was to test the impact on the quality of published reports of an intervention which would request, at the time of manuscript submission, that authors complete a checklist detailing where in the manuscript the various components of the ARRIVE checklist were met. This study, to our knowledge, is the first randomised controlled trial of requested ARRIVE guideline completion.

## Methods

### Methodology and open data

Our protocol, data analysis plan, analysis code, data validation code, and complete dataset are available on the Open Science Framework (http://dx.doi.org/10.17605/OSF.IO/XSJBV)

### Ethical approval

We sought an informal ethical opinion from the BMJ Ethics Committee, who were prepared to consider our proposal although it was slightly out of scope. We did this because we were unable at the time to identify an institutional ethics committee who considered this research to fall within their remit. The majority view of the committee was that it was ethical for manuscripts to be randomised between different handling methods; that it was ethical for authors, peer reviewers and academic editors to be kept unaware of the existence of the study while it was in progress; and that it was ethical for the study to receive funding from the NC3Rs.

### Randomisation of Manuscripts

We developed (http://doi.org/10.5281/zenodo.1188821) an online platform to support each stage of the project (https://ecrf1.clinicaltrials.ed.ac.uk/iicarus/).

The PLOS ONE editorial process involves an initial screening process, including a determination of whether a manuscript describes animal studies, whether it describes human studies (one manuscript might describe both); and categorises the area of research according to an established taxonomy. For studies reporting the use of animals, checks are carried out to ensure that appropriate institutional animal care and use committee/ethical approvals were in place, and authors of studies perceived to be at high risk – for instance those animal studies which used death as an endpoint - are contacted to provide a valid justification. Manuscripts are then allocated to an academic editor (AE), who assigns peer reviewers as appropriate

Manuscripts submitted to PLOS ONE between March and June 2015 describing *in vivo* animal research were randomised using the IICARus web platform to receive standard editorial processing (control group) or checklist completion requests (intervention group). The randomisation procedure used minimisation (weighted at 0.75) to ensure that country of origin (of the corresponding author) was balanced between groups.

On submission, authors receive an automated acknowledgement from the publisher that their submission had entered a screening phase. For manuscripts identified during screening to include *in vivo* research and which were randomised to the intervention, corresponding authors were informed in the post screening email that a completed ARRIVE checklist must be completed before the manuscript could advance through the review process. The email advised that this should include details of the page of their manuscript on which each ARRIVE item was addressed. If the PLOS editorial team did not receive a checklist, it was sent back to authors once more for completion. Manuscripts by authors who did not complete the checklist after the second contact, for any reason, were still passed to the next stage and continued in the study. The content of completed checklists were not checked against the manuscript for compliance at any stage.

### Blinded Manuscript Processing

Authors, AEs, and peer reviewers were blinded to the existence of the study. Study personnel took care, in their public comments, not to disclose details of the study or the journal at which the study was being conducted. The journal was not named in the study protocol. If authors enquired as to why their manuscript was being processed differently, they were to be advised that these differences were due to variation within the editorial team in the intensity with which they pursued efforts to improve the review process.

For studies randomised to the control group, PLOS ONE processed the manuscript according to their normal editorial processes.

Once a final decision regarding publication was made, the pre-publication materials for accepted manuscripts were collated by the PLOS editorial team. Where an ARRIVE checklist was included in the accepted materials for publication this was redacted, along with any reference in the text to the submission of a completed ARRIVE checklist. The format of manuscripts largely excluded any evidence that the manuscript was submitted to PLOS ONE. If a reference to PLOS ONE was discovered in the text by internal outcome assessors (within our research group), this was also redacted to prevent any change in behaviour which may result from external outcome assessors knowing which publisher was involved in the study. Where authors stated that the work complies with the ARRIVE guidelines this statement was not redacted. Redacted PDFs of all materials were provided to our research team and uploaded to the IICARus web platform.

### Outcome assessment

Our primary outcome was to assess whether the proportion of publications in each group considered to fully comply with all of the ARRIVE criteria was independent of group allocation.

Our secondary outcome measures were to assess whether:

- the proportion of publications meeting each of the individual 38 ARRIVE sub-items was independent of group allocation (intervention/ control)
- the proportion of studies reporting all Landis criteria (Landis et al., 2012) risk of bias items (randomisation, blinded assessment of outcome, sample size calculation and criteria for exclusion of experimental subjects) was independent of group allocation
- the proportion of submitted manuscripts accepted for publication was independent of group allocation

Our tertiary outcomes, we assessed whether:

- the proportion of publications meeting each of the 38 ARRIVE sub-items was independent of group allocation, stratified by experimental animal
- the proportion of studies reporting all of the Landis criteria risk of bias items (blinded assessment of outcome, sample size calculation and criteria for exclusion of experimental subjects), stratified by experimental animal, was independent of group allocation
- the proportion of publications meeting each of the 38 ARRIVE sub-items was independent of group allocation, stratified by the country of the address of the corresponding author
- the proportion of publications meeting each of the 38 ARRIVE sub-items was independent of group allocation, stratified by whether or not the research also contains human data

To examine the feasibility of implementing requests for ARRIVE checklist completion at PLOS ONE, we also assessed the following for accepted manuscripts in each group:

- Time (days) spent in PLOS editorial office in handling the manuscript (prior to editor assignment).
- Time (days) from manuscript submission to AE assignment.
- Time (days) from AE assignment to first reviewer agreed.
- Time (days) from AE assignment to first decision
- Time (days) from receipt of last review to AE decision
- In addition, we assessed the following outcome measures for manuscripts which were accepted following resubmission:
- Time (days) from initial decision letter to resubmission.
- Number of cycles of resubmission.
- Time (days) from resubmission to final decision.

We also conducted some exploratory analyses not defined in the study protocol to investigate:

- The proportion of publications meeting each of the 38 ARRIVE sub-items for publications in the intervention group where authors had completed an ARRIVE checklist (equivalent to an “on treatment” analysis)
- The proportion of publications meeting each Landis criteria item in each group

We operationalised the 38 subitems of the ARRIVE checklist into 108 questions which were scored by trained outcome assessors on the web platform (Appendix 1). It was later determined by the steering committee that 7 questions from the original 108 were not strictly required to comply with the ARRIVE checklist and were therefore excluded from the analysis.

PDF files of manuscripts were available alongside the scoring questions. Each manuscript was scored by two independent reviewers who were blinded to both intervention status and to the score given by the alternative reviewer. Manuscripts were presented to reviewers in random order, and the platform did not allow the same user to review the same manuscript twice. Discrepancies between reviewers were reconciled by a third reviewer, who could view both previous scores.

There were several deviations from the outcome measures specified our study protocol. The time spent in the PLOS editorial office was not disentangled from time with the authors, therefore it includes time for the authors to follow any copyediting changes and requests for documents (including the request to complete an ARRIVE checklist). Similarly, the time spent with authors was also included in the time from manuscript submission to AE assignment. In addition, we had originally intended to analyse the time in the PLOS editorial office in minutes, but the measurement of this was not feasible. We were unable to analyse “The proportion of submitted manuscripts accepted for publication, stratified by experimental animal” (Secondary outcome measure) as we did not receive species categorisation data for studies which were not accepted. PLOS ONE were unable to provide us with one of our specified feasibility measures “Time (days) for each reviewer, from solicitation of reviews to receipt of reviews)” and instead provided “Time (days) from AE assignment to first decision”. “Time (days) from AE assignment to first reviewer agreed” also differs to the outcome described in our study protocol (“Time (days) from AE assignment to solicitation of reviews”). In addition, we had originally set out in our protocol that we would look at the following feasibility outcome measures for manuscripts when the decision was other than “Accept” or “Reject”: whether a revised manuscript is submitted, time (days) from initial decision letter to resubmission, number of cycles of resubmission, time (days) from resubmission to final decision. However, we did not attain this information for manuscripts which were not eventually accepted. Therefore, all feasibility measures apply to accepted manuscripts only.

### Reviewer Training

This was a challenging project, and we used crowdsourcing to recruit additional reviewers external to our research group. We used our research networks and social media to identify researchers and students across the biomedical sciences and recruit them as outcome assessors for the project. As an incentive, rewards were given to external reviewers who reached a pre-specified number of manuscript reviews or completed the most reviews in a certain time period.

To ensure that reviewer quality was high, we required reviewers to complete online training prior to reviewing manuscripts as part of the project. We developed a training program with a pool of 10 manuscripts for which we described “Gold standard” correct answers with explanations; and an accompanying document with further elaboration (Appendix 2). To successfully complete the training, external reviewers had to score 80% against these gold standard answers overall and score 100% on gold standard questions relating to the Landis criteria items for three consecutive training papers. The training platform remains available (https://ecrf1.clinicaltrials.ed.ac.uk/iicarus/) and can still be used as a training tool for assessing manuscripts against the ARRIVE guidelines.

### Power Calculations

When the study was being designed the PLOS ONE editorial team estimated that complete compliance with the ARRIVE guidelines was close to zero. To have 80% power with an alpha of 0.05 to detect an increase in full compliance from 1% to 10% (the primary outcome) would require 100 published manuscripts per group. To examine each of the individual 38 ARRIVE subitems (Secondary outcome), after correction for multiplicity of testing (alpha = 0.0013) we would require 200 published manuscripts per group to detect with 80% power an increase from 30% to 50% in the prevalence of reporting of an individual subitem. It was estimated that at time of the trial PLOS ONE accepted around 70% of manuscripts, and to account for some drop out because of the use of the same academic editor, we increased our group estimate to 150 manuscripts per group for the primary outcome, and 300 manuscripts in each group for secondary outcomes. During the course of the study it appeared that acceptance rates were lower than the estimate, and so we increased target recruitment to 1000 manuscripts, of which we estimated 600 would be accepted for publication. We did not curtail the study when we had reached the required number of manuscripts accepted for publication because we were concerned that manuscripts with short submission to acceptance times would be enriched in the study population and might not be representative of all manuscripts.

### Data Validation

We validated the dataset, blinded to group allocation, to minimise errors. For example, where a paper was assessed as not including fish experiments, “Yes” or “No” responses to IICARus questions only relevant to fish species (e.g. Appendix 1, Questions 9.2.3 - 9.2.4), should not have been recorded. The R code for validation, with explanations of each response validated, and the changes we made to the data are available on the OSF. These were uploaded prior to the unblinding of the final results, at which point database lock occurred and the data were not subsequently altered in any way.

### Statistical Analysis

All analyses were carried out using RStudio v1.0.143 with the level of statistical significance set at p<0.05, corrected as appropriate for multiple comparisons. Our full statistical analysis plan and accompanying R code was uploaded to the OSF prior to database lock.

We performed logistic regression with group allocation and corresponding author country of origin included as independent variables to determine any effects on full compliance (Primary Outcome) and compliance with each of *the* 38 items (Secondary Outcome), adjusting for stratified randomisation. For our primary, secondary, and tertiary outcome measures we used the Chi Squared Test of Independence to test whether compliance was independent of group membership (Intervention/ Control). To determine if the differences between proportions is meaningful for each outcome measure, effect sizes were calculated using Cohen’s H. For feasibility outcomes, medians and inter-quartile ranges were calculated and the Mann-Whitney U test was used to test whether a significant difference existed between groups. To control the familywise error rate of multiple comparisons, the Holm-Bonferroni method (Aickin and Gensler, 1996) was used to adjust p-values for secondary, tertiary, and feasibility outcomes. Only means and confidence intervals were calculated for our exploratory analyses.

### Statistical Considerations

The proportion of compliant manuscripts was assessed based on the number compliant manuscripts divided by the number of applicable manuscripts. In some cases, particularly when stratifying by manuscript country of origin or animal species, the number of manuscripts in each group is very low. If the number of applicable manuscripts for any subitem (with or without stratification) was less than 10, we did not perform statistical analysis.

The Chi Squared Test of Independence relies on the assumption that no more than 20% of expected counts are less than 5 and that no individual expected counts are less than 1. In cases where counts were less than 5, a Fisher’s exact test was used.

Unpaired t-tests rely on a normal distribution, therefore if the distribution was non-normal the Mann-Whitney U test, a non-parametric alternative, was used and summary medians and inter-quartile ranges were presented. In the case of parametric data with unequal variance between groups Welch’s t-test was used due to higher reliability.

## Results

We randomised 1689 PLOS ONE manuscripts; 845 manuscripts to the intervention and 844 to control. We later excluded 420 manuscripts which in retrospect did not meet the inclusion criteria (largely because they described *ex vivo* rather than *in vivo* research). Of the remaining 1269 manuscripts, 672 were accepted for publication (340 control, 332 intervention) and underwent web based outcome assessment (Figure 1). Manuscript allocation to group, and the corresponding number of manuscripts from each country post-randomisation and post-acceptance is shown in Table 1. No authors questioned the differences in manuscript processing occurring within the intervention group. A complete dataset detailing the proportion compliance for each of the 108 questions is available online (http://dx.doi.org/10.17605/OSF.IO/XSJBV).

**Table 1:**
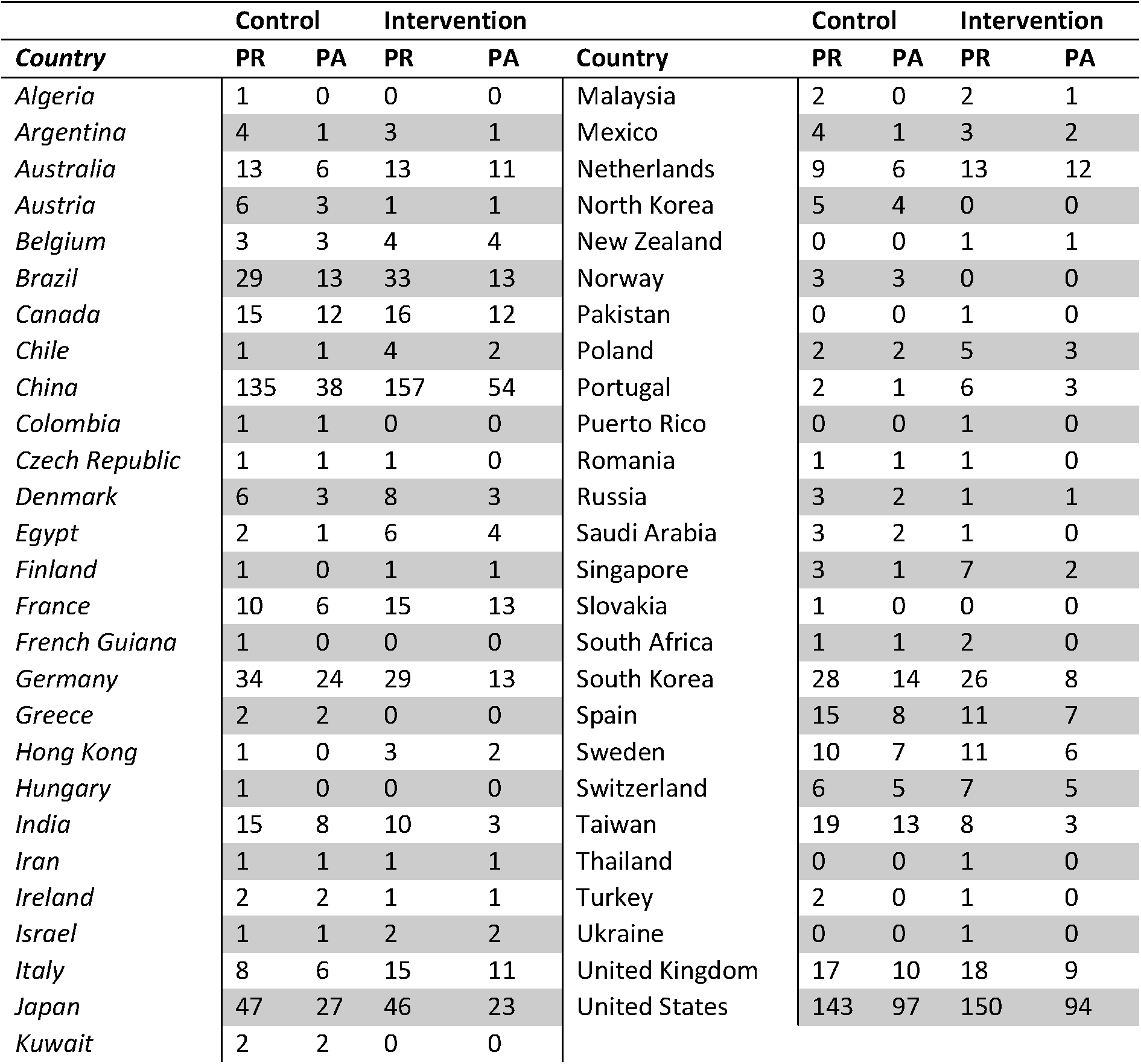
Manuscript allocation by country; Manuscripts allocated to each group per corresponding author country of origin; PR, post randomisation; PA, post-acceptance

|  |  | Control |  | Intervention |  |  |  | Control |  | Intervention |  |
| --- | --- | --- | --- | --- | --- | --- | --- | --- | --- | --- | --- |
| *Country* |  | PR | PA | PR | PA | *Country* |  | PR | PA | PR | PA |
| *Algeria* |  | 1 | 0 | 0 | 0 | *Malaysia* |  | 2 | 0 | 2 | 1 |
| *Argentina* |  | 4 | 1 | 3 | 1 | *Mexico* |  | 4 | 1 | 3 | 2 |
| *Australia* |  | 13 | 6 | 13 | 11 | *Netherlands* |  | 9 | 6 | 13 | 12 |
| *Austria* |  | 6 | 3 | 1 | 1 | *North Korea* |  | 5 | 4 | 0 | 0 |
| *Belgium* |  | 3 | 3 | 4 | 4 | *New Zealand* |  | 0 | 0 | 1 | 1 |
| *Brazil* |  | 29 | 13 | 33 | 13 | *Norway* |  | 3 | 3 | 0 | 0 |
| *Canada* |  | 15 | 12 | 16 | 12 | *Pakistan* |  | 0 | 0 | 1 | 0 |
| *Chile* |  | 1 | 1 | 4 | 2 | *Poland* |  | 2 | 2 | 5 | 3 |
| *China* |  | 135 | 38 | 157 | 54 | *Portugal* |  | 2 | 1 | 6 | 3 |
| *Colombia* |  | 1 | 1 | 0 | 0 | *Puerto Rico* |  | 0 | 0 | 1 | 0 |
| *Czech Republic* |  | 1 | 1 | 1 | 0 | *Romania* |  | 1 | 1 | 1 | 0 |
| *Denmark* |  | 6 | 3 | 8 | 3 | *Russia* |  | 3 | 2 | 1 | 1 |
| *Egypt* |  | 2 | 1 | 6 | 4 | *Saudi Arabia* |  | 3 | 2 | 1 | 0 |
| *Finland* |  | 1 | 0 | 1 | 1 | *Singapore* |  | 3 | 1 | 7 | 2 |
| *France* |  | 10 | 6 | 15 | 13 | *Slovakia* |  | 1 | 0 | 0 | 0 |
| *French Guiana* |  | 1 | 0 | 0 | 0 | *South Africa* |  | 1 | 1 | 2 | 0 |
| *Germany* |  | 34 | 24 | 29 | 13 | *South Korea* |  | 28 | 14 | 26 | 8 |
| *Greece* |  | 2 | 2 | 0 | 0 | *Spain* |  | 15 | 8 | 11 | 7 |
| *Hong Kong* |  | 1 | 0 | 3 | 2 | *Sweden* |  | 10 | 7 | 11 | 6 |
| *Hungary* |  | 1 | 0 | 0 | 0 | *Switzerland* |  | 6 | 5 | 7 | 5 |
| *India* |  | 15 | 8 | 10 | 3 | *Taiwan* |  | 19 | 13 | 8 | 3 |
| *Iran* |  | 1 | 1 | 1 | 1 | *Thailand* |  | 0 | 0 | 1 | 0 |
| *Ireland* |  | 2 | 2 | 1 | 1 | *Turkey* |  | 2 | 0 | 1 | 0 |
| *Israel* |  | 1 | 1 | 2 | 2 | *Ukraine* |  | 0 | 0 | 1 | 0 |
| *Italy* |  | 8 | 6 | 15 | 11 | *United Kingdom* |  | 17 | 10 | 18 | 9 |
| *Japan* |  | 47 | 27 | 46 | 23 | *United States* |  | 143 | 97 | 150 | 94 |
| *Kuwait* |  | 2 | 2 | 0 | 0 |  |  |  |  |  |  |

**Table 2:** Percentage compliance for each ARRIVE subitem; %, percentage of compliant papers; CI, confidence interval; n, number of compliant papers; N, total number of applicable papers; Adj p, adjusted p value; n.s, not significant; Cohen’s H, Cohen’s H effect size

| *ARRIVE Item* | Control |  |  |  | Intervention |  |  |  | Adj. p | Cohen's H |
| --- | --- | --- | --- | --- | --- | --- | --- | --- | --- | --- |
|  | % | 95% CIs | n | N | % | 95% CIs | n | N |  |  |
| 1 | 41.76 | 36.5-47.2 | 142 | 340 | 44.58 | 39.2-50.1 | 148 | 332 | n.s. | 0.06 |
| 2 | 71.76 | 66.6-76.4 | 244 | 340 | 67.47 | 62.1-72.4 | 224 | 332 | n.s. | -0.09 |
| 3a | 100.00 | 98.6-100 | 340 | 340 | 100.00 | 98.6-100 | 332 | 332 | n.s. | 0.00 |
| 3b | 34.12 | 29.1-39.5 | 116 | 340 | 36.14 | 31-41.6 | 120 | 332 | n.s. | 0.04 |
| 4 | 91.18 | 87.5-93.9 | 310 | 340 | 93.07 | 89.6-95.5 | 309 | 332 | n.s. | 0.07 |
| 5 | 69.41 | 64.2-74.2 | 236 | 340 | 72.59 | 67.4-77.3 | 241 | 332 | n.s. | 0.07 |
| 6a | 70.00 | 64.8-74.8 | 238 | 340 | 75.00 | 69.9-79.5 | 249 | 332 | n.s. | 0.11 |
| 6b | 8.33 | 5.7-12 | 28 | 336 | 10.49 | 7.5-14.5 | 34 | 324 | n.s. | 0.07 |
| 6c | 90.00 | 86.2-92.9 | 306 | 340 | 88.86 | 84.8-91.9 | 295 | 332 | n.s. | -0.04 |
| 7a | 16.76 | 13-21.3 | 57 | 340 | 16.87 | 13.1-21.4 | 56 | 332 | n.s. | 0.00 |
| 7b | 44.37 | 38.7-50.2 | 134 | 302 | 51.33 | 45.5-57.1 | 154 | 300 | n.s. | 0.14 |
| 7c | 8.64 | 5.8-12.5 | 26 | 301 | 14.09 | 10.5-18.7 | 42 | 298 | n.s. | 0.17 |
| 7d | 3.63 | 1.9-6.6 | 11 | 303 | 3.63 | 1.9-6.6 | 11 | 303 | n.s. | 0.00 |
| 8a | 4.71 | 2.8-7.7 | 16 | 340 | 7.83 | 5.3-11.4 | 26 | 332 | n.s. | 0.13 |
| 8b | 57.06 | 51.6-62.4 | 194 | 340 | 62.65 | 57.2-67.8 | 208 | 332 | n.s. | 0.11 |
| 9a | 0.30 | 0-1.9 | 1 | 337 | 2.74 | 1.3-5.3 | 9 | 328 | n.s. | 0.22 |
| 9b | 52.06 | 46.6-57.5 | 177 | 340 | 74.10 | 69-78.7 | 246 | 332 | <0.001 | 0.46 |
| 9c | 14.71 | 11.2-19 | 50 | 340 | 20.48 | 16.4-25.3 | 68 | 332 | n.s. | 0.15 |
| 10a | 37.35 | 32.2-42.8 | 127 | 340 | 43.67 | 38.3-49.2 | 145 | 332 | n.s. | 0.13 |
| 10b | 3.53 | 1.9-6.2 | 12 | 340 | 7.53 | 5-11.1 | 25 | 332 | n.s. | 0.18 |
| 10c | 18.15 | 14.3-22.8 | 61 | 336 | 14.64 | 11.1-19.1 | 47 | 321 | n.s. | -0.10 |
| 11a | 4.82 | 2.8-8 | 15 | 311 | 7.49 | 4.9-11.2 | 23 | 307 | n.s. | 0.11 |
| 11b | 1.24 | 0.4-3.4 | 4 | 323 | 2.88 | 1.4-5.6 | 9 | 313 | n.s. | 0.12 |
| 12 | 1.76 | 0.7-4 | 6 | 340 | 3.01 | 1.5-5.6 | 10 | 332 | n.s. | 0.08 |
| 13a | 87.50 | 83.4-90.7 | 294 | 336 | 89.91 | 86-92.9 | 294 | 327 | n.s. | 0.08 |
| 13b | 44.08 | 38.7-49.6 | 149 | 338 | 44.51 | 39.1-50.1 | 146 | 328 | n.s. | 0.01 |
| 13c | 10.06 | 7.2-13.9 | 34 | 338 | 12.80 | 9.5-17 | 42 | 328 | n.s. | 0.09 |
| 14 | 0.29 | 0-1.9 | 1 | 340 | 0.00 | 0-1.4 | 0 | 332 | n.s. | -0.11 |
| 15a | 37.35 | 32.2-42.8 | 127 | 340 | 37.35 | 32.2-42.8 | 124 | 332 | n.s. | 0.00 |
| 15b | 12.65 | 9.4-16.8 | 43 | 340 | 14.46 | 10.9-18.8 | 48 | 332 | n.s. | 0.05 |
| 16 | 78.55 | 73.7-82.8 | 260 | 331 | 80.94 | 76.1-85 | 259 | 320 | n.s. | 0.06 |
| 17a | 16.47 | 12.8-20.9 | 56 | 340 | 21.69 | 17.5-26.6 | 72 | 332 | n.s. | 0.13 |
| 17b | 1.18 | 0.4-3.2 | 4 | 340 | 1.81 | 0.7-4.1 | 6 | 332 | n.s. | 0.05 |
| 18a | 100.00 | 98.6-100 | 340 | 340 | 99.40 | 97.6-99.9 | 330 | 332 | n.s. | -0.16 |
| 18b | 26.47 | 21.9-31.6 | 90 | 340 | 28.31 | 23.6-33.5 | 94 | 332 | n.s. | 0.04 |
| 18c | 2.94 | 1.5-5.5 | 10 | 340 | 3.01 | 1.5-5.6 | 10 | 332 | n.s. | 0.00 |
| 19 | 77.94 | 73.1-82.2 | 265 | 340 | 77.71 | 72.8-82 | 258 | 332 | n.s. | -0.01 |
| 20 | 51.47 | 46-56.9 | 175 | 340 | 52.71 | 47.2-58.2 | 175 | 332 | n.s. | 0.02 |

**Figure 1:**
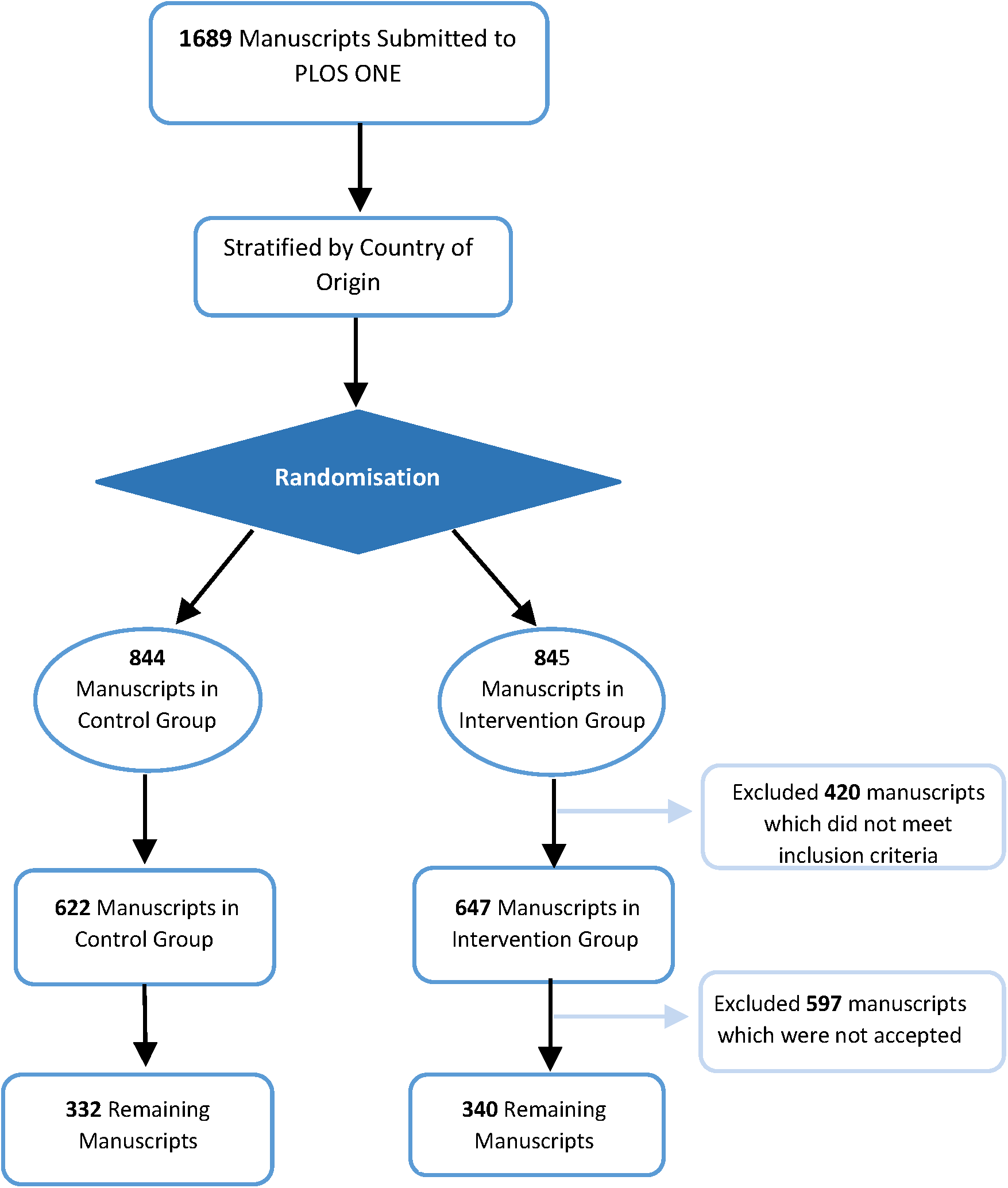
Manuscript processing

### Quality of Outcome Assessment

360 individuals registered with the online platform; 47 completed reviewer training and 42 contributed at least one outcome assessment. The percentage agreement between the first and second reviewer for each manuscript was high. For the majority (71.6%) of manuscripts, reviewers were in agreement on at least 80% of the questions. The agreement of reviewers varied considerably at the level of each of the 108 individual questions (Supplementary Table 1), from a kappa coefficient of 0.90 (0.86 - 0.93) for Question 1.1 *(Is the species of animal model studied reported in the title?*) to a worse than chance kappa coefficient of −0.03 (−0.10 - −0.04) for Question 13.2 *(Is the unit of analysis for at least one test explicitly specified?)*. This distribution of kappa agreement is displayed in a histogram (Supplementary Figure 1)

### Primary Outcome

No manuscript achieved full compliance with the ARRIVE checklist, therefore there was no difference between intervention and control groups. Compliance with individual ARRIVE items ranged from 8% to 65%. The median compliance was 36.8 % and 39.5% of relevant items in the control and intervention groups respectively.

### Secondary Outcomes

#### Logistic Regression

Manuscript country of corresponding author had no influence on compliance either overall or for any individual subitems. Only one subitem had improved reporting in the intervention group, subitem 9b (*Provide details of husbandry conditions e.g. breeding programme, light/dark cycle, temperature, quality of water etc for fish, type of food, access to food and water, environmental enrichment*) (increased log odds of compliance by 1.03 (p<0.0001).

#### Compliance with individual ARRIVE Subitems

Only one ARRIVE item had improved compliance in the intervention group. Reporting of ARRIVE subitem, 9b increased significantly from 52.1% (177/340) in the control group to 74.1% (246/332) in the intervention group (X^2^ = 34.0, df=1, p<0.0001). Reporting of animal characteristics and health status (Item 14) was very low, with 0.29% (1/339) and 0% (0/332) compliance in the control and intervention groups respectively. Similarly, reporting of animal housing (Item 9a), adverse events (Item 17b), the order of treatment and assessment (Item 11b), implications for replacement, refinement, or reduction (Item 18c), defining primary and secondary outcomes (Item 12), and rationale for experimental procedures (Item 7d) was low, with less than 5% of manuscripts reporting each of these items in both groups. Figure 2 shows the percentage compliance in each group for each ARRIVE subitem in each section of the manuscript.

**Figure 2:**
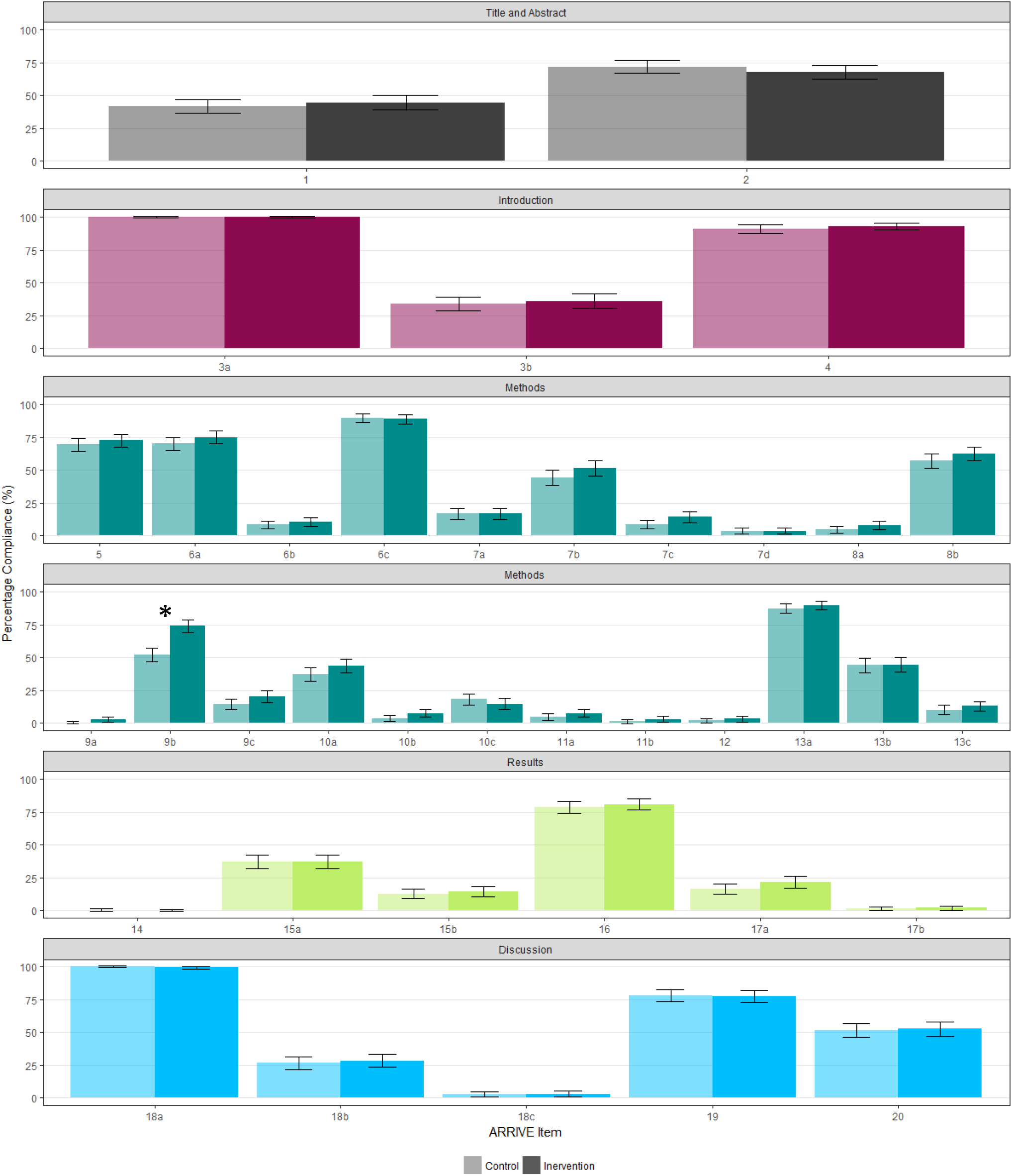
Percentage compliance for each ARRIVE subitem; Percentage compliance for each ARRIVE item with 95% confidence intervals; * denotes significance; figure divided into article sections specified in the ARRIVE guidelines

#### Reporting of Landis 4 items

Reporting of the Landis 4 criteria (blinding, randomisation, animal exclusions, and use of a sample size calculation) was low and did not differ between groups (X^2^ = 16.8, df=1, p=0.003). 1.5% of the control group manuscripts (5/340) and 0.9% (3/332) of intervention group manuscripts reported all 4 items of the Landis criteria (Fisher’s estimate for difference = 0.61, df=1, p=0.73).

#### Manuscript Acceptance

There was no difference in the proportion of accepted manuscripts between the control and intervention groups, being 54.7% (340/622) and 51.3% (322/647) respectively.

### Tertiary Outcomes

#### Compliance by Animal Species

In studies involving mice, reporting of one ARRIVE subitem, 9b (husbandry related) increased significantly from 49.5% (105/211) in the control group to 70.2% (135/192) in the intervention group (X^2^ = 16.8, df=1, p=0.003). No subitem had significant differences between groups in rat studies. Results are summarised in Table 3a and Table 3b. There was no difference in Landis 4 compliance between animal species.

**Table 3:**
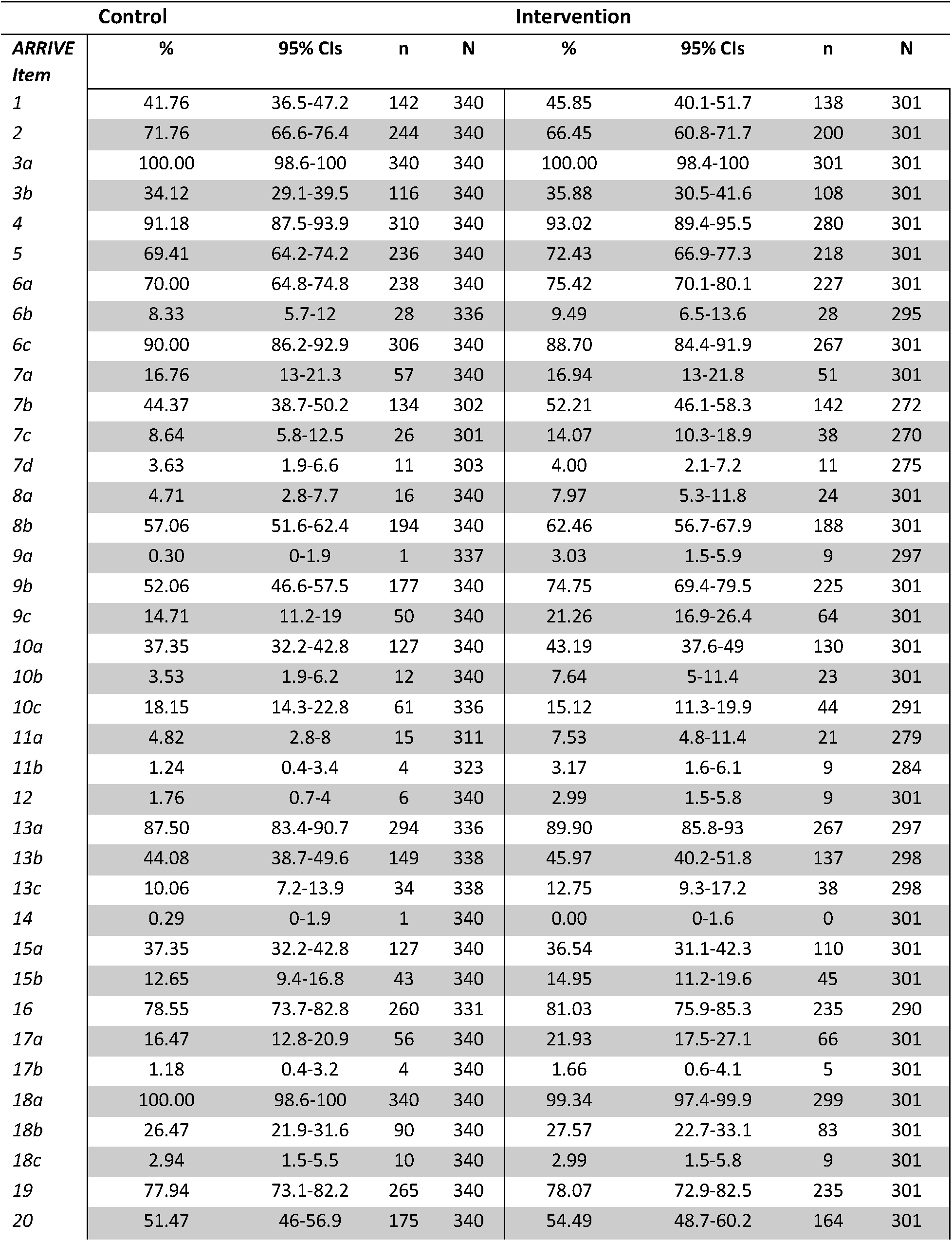
True intervention ARRIVE subitem compliance; %, percentage of compliant papers; CI, confidence interval; n, number of compliant papers; N, total number of applicable papers; Adj p, adjusted p value; n.s, not significant; Cohen’s H, Cohen’s H effect size

| *ARRIVE Item* | Control |  |  |  | Intervention |  |  |  |
| --- | --- | --- | --- | --- | --- | --- | --- | --- |
|  | % | 95% CIs | n | N | % | 95% CIs | n | N |
| 1 | 41.76 | 36.5-47.2 | 142 | 340 | 45.85 | 40.1-51.7 | 138 | 301 |
| 2 | 71.76 | 66.6-76.4 | 244 | 340 | 66.45 | 60.8-71.7 | 200 | 301 |
| 3a | 100.00 | 98.6-100 | 340 | 340 | 100.00 | 98.4-100 | 301 | 301 |
| 3b | 34.12 | 29.1-39.5 | 116 | 340 | 35.88 | 30.5-41.6 | 108 | 301 |
| 4 | 91.18 | 87.5-93.9 | 310 | 340 | 93.02 | 89.4-95.5 | 280 | 301 |
| 5 | 69.41 | 64.2-74.2 | 236 | 340 | 72.43 | 66.9-77.3 | 218 | 301 |
| 6a | 70.00 | 64.8-74.8 | 238 | 340 | 75.42 | 70.1-80.1 | 227 | 301 |
| 6b | 8.33 | 5.7-12 | 28 | 336 | 9.49 | 6.5-13.6 | 28 | 295 |
| 6c | 90.00 | 86.2-92.9 | 306 | 340 | 88.70 | 84.4-91.9 | 267 | 301 |
| 7a | 16.76 | 13-21.3 | 57 | 340 | 16.94 | 13-21.8 | 51 | 301 |
| 7b | 44.37 | 38.7-50.2 | 134 | 302 | 52.21 | 46.1-58.3 | 142 | 272 |
| 7c | 8.64 | 5.8-12.5 | 26 | 301 | 14.07 | 10.3-18.9 | 38 | 270 |
| 7d | 3.63 | 1.9-6.6 | 11 | 303 | 4.00 | 2.1-7.2 | 11 | 275 |
| 8a | 4.71 | 2.8-7.7 | 16 | 340 | 7.97 | 5.3-11.8 | 24 | 301 |
| 8b | 57.06 | 51.6-62.4 | 194 | 340 | 62.46 | 56.7-67.9 | 188 | 301 |
| 9a | 0.30 | 0-1.9 | 1 | 337 | 3.03 | 1.5-5.9 | 9 | 297 |
| 9b | 52.06 | 46.6-57.5 | 177 | 340 | 74.75 | 69.4-79.5 | 225 | 301 |
| 9c | 14.71 | 11.2-19 | 50 | 340 | 21.26 | 16.9-26.4 | 64 | 301 |
| 10a | 37.35 | 32.2-42.8 | 127 | 340 | 43.19 | 37.6-49 | 130 | 301 |
| 10b | 3.53 | 1.9-6.2 | 12 | 340 | 7.64 | 5-11.4 | 23 | 301 |
| 10c | 18.15 | 14.3-22.8 | 61 | 336 | 15.12 | 11.3-19.9 | 44 | 291 |
| 11a | 4.82 | 2.8-8 | 15 | 311 | 7.53 | 4.8-11.4 | 21 | 279 |
| 11b | 1.24 | 0.4-3.4 | 4 | 323 | 3.17 | 1.6-6.1 | 9 | 284 |
| 12 | 1.76 | 0.7-4 | 6 | 340 | 2.99 | 1.5-5.8 | 9 | 301 |
| 13a | 87.50 | 83.4-90.7 | 294 | 336 | 89.90 | 85.8-93 | 267 | 297 |
| 13b | 44.08 | 38.7-49.6 | 149 | 338 | 45.97 | 40.2-51.8 | 137 | 298 |
| 13c | 10.06 | 7.2-13.9 | 34 | 338 | 12.75 | 9.3-17.2 | 38 | 298 |
| 14 | 0.29 | 0-1.9 | 1 | 340 | 0.00 | 0-1.6 | 0 | 301 |
| 15a | 37.35 | 32.2-42.8 | 127 | 340 | 36.54 | 31.1-42.3 | 110 | 301 |
| 15b | 12.65 | 9.4-16.8 | 43 | 340 | 14.95 | 11.2-19.6 | 45 | 301 |
| 16 | 78.55 | 73.7-82.8 | 260 | 331 | 81.03 | 75.9-85.3 | 235 | 290 |
| 17a | 16.47 | 12.8-20.9 | 56 | 340 | 21.93 | 17.5-27.1 | 66 | 301 |
| 17b | 1.18 | 0.4-3.2 | 4 | 340 | 1.66 | 0.6-4.1 | 5 | 301 |
| 18a | 100.00 | 98.6-100 | 340 | 340 | 99.34 | 97.4-99.9 | 299 | 301 |
| 18b | 26.47 | 21.9-31.6 | 90 | 340 | 27.57 | 22.7-33.1 | 83 | 301 |
| 18c | 2.94 | 1.5-5.5 | 10 | 340 | 2.99 | 1.5-5.8 | 9 | 301 |
| 19 | 77.94 | 73.1-82.2 | 265 | 340 | 78.07 | 72.9-82.5 | 235 | 301 |
| 20 | 51.47 | 46-56.9 | 175 | 340 | 54.49 | 48.7-60.2 | 164 | 301 |

**Table 3a:** ARRIVE item compliance in mouse studies. %, percentage of compliant papers; CI, confidence interval; n, number of compliant papers; N, total number of applicable papers; Adj p, adjusted p value; n.s, not significant; Cohen’s H, Cohen’s H effect size

|  | Control |  |  |  | Intervention |  |  |  |  |  |
| --- | --- | --- | --- | --- | --- | --- | --- | --- | --- | --- |
| *ARRIVE Item* | % | 95% CIs | n | N | % | 95% CIs | n | N | Adj p | Cohen's H |
| 1 | 37.44 | 31-44.4 | 79 | 211 | 37.50 | 30.7-44.8 | 72 | 192 | n.s. | 0.00 |
| 2 | 68.25 | 61.4-74.4 | 144 | 211 | 60.42 | 53.1-67.3 | 116 | 192 | n.s. | -0.16 |
| 3a | 100.00 | 97.8-100 | 211 | 211 | 100.00 | 97.6-100 | 192 | 192 | n.s. | 0.00 |
| 3b | 30.33 | 24.3-37.1 | 64 | 211 | 29.69 | 23.4-36.8 | 57 | 192 | n.s. | -0.01 |
| 4 | 89.10 | 83.9-92.8 | 188 | 211 | 92.19 | 87.2-95.4 | 177 | 192 | n.s. | 0.11 |
| 5 | 67.77 | 61-73.9 | 143 | 211 | 71.88 | 64.9-78 | 138 | 192 | n.s. | 0.09 |
| 6a | 63.98 | 57.1-70.4 | 135 | 211 | 71.35 | 64.3-77.5 | 137 | 192 | n.s. | 0.16 |
| 6b | 5.24 | 2.8-9.4 | 11 | 210 | 7.89 | 4.6-12.9 | 15 | 190 | n.s. | 0.11 |
| 6c | 90.05 | 85-93.6 | 190 | 211 | 91.15 | 86-94.6 | 175 | 192 | n.s. | 0.04 |
| 7a | 17.54 | 12.8-23.5 | 37 | 211 | 17.71 | 12.7-24 | 34 | 192 | n.s. | 0.00 |
| 7b | 44.68 | 37.5-52.1 | 84 | 188 | 48.57 | 41-56.2 | 85 | 175 | n.s. | 0.08 |
| 7c | 6.91 | 3.9-11.8 | 13 | 188 | 9.83 | 6-15.5 | 17 | 173 | n.s. | 0.11 |
| 7d | 3.16 | 1.3-7.1 | 6 | 190 | 2.27 | 0.7-6.1 | 4 | 176 | n.s. | -0.05 |
| 8a | 4.74 | 2.4-8.8 | 10 | 211 | 6.25 | 3.4-10.9 | 12 | 192 | n.s. | 0.07 |
| 8b | 55.45 | 48.5-62.2 | 117 | 211 | 60.94 | 53.6-67.8 | 117 | 192 | n.s. | 0.11 |
| 9a | 0.47 | 0-3 | 1 | 211 | 2.08 | 0.7-5.6 | 4 | 192 | n.s. | 0.15 |
| 9b | 49.76 | 42.8-56.7 | 105 | 211 | 70.31 | 63.2-76.6 | 135 | 192 | 0.003 | 0.42 |
| 9c | 15.17 | 10.7-20.9 | 32 | 211 | 23.96 | 18.2-30.7 | 46 | 192 | n.s. | 0.22 |
| 10a | 27.01 | 21.3-33.6 | 57 | 211 | 31.25 | 24.9-38.4 | 60 | 192 | n.s. | 0.09 |
| 10b | 2.84 | 1.2-6.4 | 6 | 211 | 5.73 | 3-10.3 | 11 | 192 | n.s. | 0.14 |
| 10c | 21.15 | 15.9-27.5 | 44 | 208 | 15.43 | 10.7-21.6 | 29 | 188 | n.s. | -0.15 |
| 11a | 4.12 | 1.9-8.3 | 8 | 194 | 5.00 | 2.5-9.6 | 9 | 180 | n.s. | 0.04 |
| 11b | 1.93 | 0.6-5.2 | 4 | 207 | 0.54 | 0-3.4 | 1 | 185 | n.s. | -0.13 |
| 12 | 0.00 | 0-2.2 | 0 | 211 | 3.65 | 1.6-7.7 | 7 | 192 | n.s. | 0.38 |
| 13a | 87.20 | 81.8-91.3 | 184 | 211 | 89.58 | 84.2-93.4 | 172 | 192 | n.s. | 0.07 |
| 13b | 46.92 | 40.1-53.9 | 99 | 211 | 42.71 | 35.7-50 | 82 | 192 | n.s. | -0.08 |
| 13c | 8.06 | 4.9-12.8 | 17 | 211 | 11.46 | 7.5-17 | 22 | 192 | n.s. | 0.12 |
| 14 | 0.00 | 0-2.2 | 0 | 211 | 0.00 | 0-2.4 | 0 | 192 | n.s. | 0.00 |
| 15a | 36.02 | 29.6-42.9 | 76 | 211 | 36.46 | 29.7-43.7 | 70 | 192 | n.s. | 0.01 |
| 15b | 11.37 | 7.6-16.6 | 24 | 211 | 10.94 | 7.1-16.4 | 21 | 192 | n.s. | -0.01 |
| 16 | 84.62 | 78.8-89.1 | 176 | 208 | 80.95 | 74.5-86.1 | 153 | 189 | n.s. | -0.10 |
| 17a | 15.64 | 11.2-21.4 | 33 | 211 | 23.44 | 17.8-30.2 | 45 | 192 | n.s. | 0.20 |
| 17b | 0.95 | 0.2-3.7 | 2 | 211 | 2.60 | 1-6.3 | 5 | 192 | n.s. | 0.13 |
| 18a | 100.00 | 97.8-100 | 211 | 211 | 98.96 | 95.9-99.8 | 190 | 192 | n.s. | -0.20 |
| 18b | 27.01 | 21.3-33.6 | 57 | 211 | 26.56 | 20.6-33.5 | 51 | 192 | n.s. | -0.01 |
| 18c | 3.32 | 1.5-7 | 7 | 211 | 3.13 | 1.3-7 | 6 | 192 | n.s. | -0.01 |
| 19 | 82.46 | 76.5-87.2 | 174 | 211 | 81.77 | 75.4-86.8 | 157 | 192 | n.s. | -0.02 |
| 20 | 51.18 | 44.2-58.1 | 108 | 211 | 56.25 | 48.9-63.3 | 108 | 192 | n.s. | 0.10 |

**Table 3b:** ARRIVE item compliance in rat studies %, percentage of compliant papers; CI, confidence interval; n, number of compliant papers; N, total number of applicable papers; Adj p, adjusted p value; n.s, not significant; Cohen’s H, Cohen’s H effect size

|  | Control |  |  |  | Intervention |  |  |  |  |  |
| --- | --- | --- | --- | --- | --- | --- | --- | --- | --- | --- |
| *ARRIVE Item* | % | 95% CIs | n | N | % | 95% CIs | n | N | Adj p | Cohen's H |
| **1** | 47.27 | 33.9-61.1 | 26 | 55 | 59.09 | 46.3-70.8 | 39 | 66 | n.s. | 0.24 |
| **2** | 83.64 | 70.7-91.8 | 46 | 55 | 81.82 | 70-89.9 | 54 | 66 | n.s. | -0.05 |
| **3a** | 100.00 | 91.9-100 | 55 | 55 | 100.00 | 93.1-100 | 66 | 66 | n.s. | 0.00 |
| **3b** | 20.00 | 10.9-33.4 | 11 | 55 | 34.85 | 23.8-47.7 | 23 | 66 | n.s. | 0.34 |
| **4** | 96.36 | 86.4-99.4 | 53 | 55 | 96.97 | 88.5-99.5 | 64 | 66 | n.s. | 0.03 |
| **5** | 83.64 | 70.7-91.8 | 46 | 55 | 80.30 | 68.3-88.7 | 53 | 66 | n.s. | -0.09 |
| **6a** | 90.91 | 79.3-96.6 | 50 | 55 | 87.88 | 77-94.3 | 58 | 66 | n.s. | -0.10 |
| **6b** | 24.53 | 14.2-38.6 | 13 | 53 | 21.54 | 12.7-33.8 | 14 | 65 | n.s. | -0.07 |
| **6c** | 98.18 | 89-99.9 | 54 | 55 | 89.39 | 78.8-95.3 | 59 | 66 | n.s. | -0.39 |
| **7a** | 9.09 | 3.4-20.7 | 5 | 55 | 12.12 | 5.7-23 | 8 | 66 | n.s. | 0.10 |
| **7b** | 44.44 | 31.2-58.5 | 24 | 54 | 60.32 | 47.2-72.2 | 38 | 63 | n.s. | 0.32 |
| **7c** | 5.56 | 1.4-16.3 | 3 | 54 | 14.29 | 7.1-25.9 | 9 | 63 | n.s. | 0.30 |
| **7d** | 1.85 | 0.1-11.2 | 1 | 54 | 6.35 | 2.1-16.3 | 4 | 63 | n.s. | 0.24 |
| **8a** | 1.82 | 0.1-11 | 1 | 55 | 10.61 | 4.7-21.2 | 7 | 66 | n.s. | 0.39 |
| **8b** | 63.64 | 49.5-75.9 | 35 | 55 | 75.76 | 63.4-85.1 | 50 | 66 | n.s. | 0.26 |
| **9a** | 0.00 | 0-8.1 | 0 | 55 | 6.15 | 2-15.8 | 4 | 65 | n.s. | 0.50 |
| **9b** | 69.09 | 55-80.5 | 38 | 55 | 90.91 | 80.6-96.3 | 60 | 66 | n.s. | 0.57 |
| **9c** | 12.73 | 5.7-25.1 | 7 | 55 | 13.64 | 6.8-24.8 | 9 | 66 | n.s. | 0.03 |
| **10a** | 52.73 | 38.9-66.1 | 29 | 55 | 60.61 | 47.8-72.2 | 40 | 66 | n.s. | 0.16 |
| **10b** | 1.82 | 0.1-11 | 1 | 55 | 12.12 | 5.7-23 | 8 | 66 | n.s. | 0.44 |
| **10c** | 5.45 | 1.4-16.1 | 3 | 55 | 3.17 | 0.6-12 | 2 | 63 | n.s. | -0.11 |
| **11a** | 3.77 | 0.7-14.1 | 2 | 53 | 10.77 | 4.8-21.5 | 7 | 65 | n.s. | 0.28 |
| **11b** | 0.00 | 0-8.4 | 0 | 53 | 4.69 | 1.2-14 | 3 | 64 | n.s. | 0.44 |
| **12** | 1.82 | 0.1-11 | 1 | 55 | 1.52 | 0.1-9.3 | 1 | 66 | n.s. | -0.02 |
| **13a** | 94.55 | 83.9-98.6 | 52 | 55 | 90.77 | 80.3-96.2 | 59 | 65 | n.s. | -0.15 |
| **13b** | 41.82 | 28.9-55.9 | 23 | 55 | 48.48 | 36.1-61 | 32 | 66 | n.s. | 0.13 |
| **13c** | 18.18 | 9.5-31.4 | 10 | 55 | 16.67 | 9-28.3 | 11 | 66 | n.s. | -0.04 |
| **14** | 0.00 | 0-8.1 | 0 | 55 | 0.00 | 0-6.9 | 0 | 66 | n.s. | 0.00 |
| **15a** | 43.64 | 30.6-57.6 | 24 | 55 | 43.94 | 31.9-56.7 | 29 | 66 | n.s. | 0.01 |
| **15b** | 12.73 | 5.7-25.1 | 7 | 55 | 16.67 | 9-28.3 | 11 | 66 | n.s. | 0.11 |
| **16** | 70.37 | 56.2-81.6 | 38 | 54 | 87.30 | 76-94 | 55 | 63 | n.s. | 0.42 |
| **17a** | 12.73 | 5.7-25.1 | 7 | 55 | 12.12 | 5.7-23 | 8 | 66 | n.s. | -0.02 |
| **17b** | 1.82 | 0.1-11 | 1 | 55 | 0.00 | 0-6.9 | 0 | 66 | n.s. | -0.27 |
| **18a** | 100.00 | 91.9-100 | 55 | 55 | 100.00 | 93.1-100 | 66 | 66 | n.s. | 0.00 |
| **18b** | 18.18 | 9.5-31.4 | 10 | 55 | 28.79 | 18.6-41.4 | 19 | 66 | n.s. | 0.25 |
| **18c** | 3.64 | 0.6-13.6 | 2 | 55 | 1.52 | 0.1-9.3 | 1 | 66 | n.s. | -0.14 |
| **19** | 74.55 | 60.7-84.9 | 41 | 55 | 86.36 | 75.2-93.2 | 57 | 66 | n.s. | 0.30 |
| **20** | 43.64 | 30.6-57.6 | 24 | 55 | 39.39 | 27.8-52.2 | 26 | 66 | n.s. | -0.09 |

#### Feasibility Measures

Re-assignment of academic editors occurred in a small number of cases (7/672), which confounds the recorded time in each stage and prevented us from analysing the feasibility outcomes for these manuscripts. The time from receipt of last review to final AE decision was missing from a significant proportion of the remaining manuscripts (342/665) and so this analysis was not performed. 10 additional manuscripts were also excluded from the feasibility dataset due to missing data on one or more feasibility outcome measures. After these exclusions, the feasibility analysis was performed on 328/340 manuscripts in the control group and 327/332 in the intervention group. For analysis of resubmitted articles, 7 manuscripts were removed as these were accepted at first decision leaving 323/340 in the control group and 325/332 in the intervention group.

Data were skewed so we used Mann-Whitney U test to compare timings between groups. Time spent in the PLOS editorial office was significantly higher (p<0.0001) for manuscripts in the intervention group with a median of 9 days (IQR=6-16.5) compared to the control group with a median of 6 days (3-10). Time from submission to academic editor assignment was also significantly higher in the intervention group (13 days, range 9-22) than in the control group (9 days, range 7-14) (p<0.0001). No significant differences were identified for other feasibility outcomes (Table 4).

**Table 4:** Feasibility measures; Q1-Q3, interquartile range; N, number of applicable manuscripts; Adj p, adjusted p value; n.s, not significant;

| Feasibility Outcomes | Control |  |  | Intervention |  |  | Adj p |
| --- | --- | --- | --- | --- | --- | --- | --- |
|  | Median | Q1 - Q3 | N | Median | Q1 - Q3 | N |  |
| Days in PLOS editorial office | 6 | 3-10 | 328 | 9 | 6-16.5 | 327 | <0.0001 |
| Days from submission to AE assignment | 9 | 7-14 | 328 | 13 | 9-22 | 327 | <0.0001 |
| Days from AE assignment to reviewer assignment | 3 | 1-8 | 328 | 3 | 1-9 | 327 | n.s. |
| Days from AE assignment first decision | 28 | 20-41.3 | 328 | 27 | 19-41 | 327 | n.s. |
| Days from initial decision to resubmission | 41 | 23.5-51.5 | 323 | 40 | 23-45 | 325 | n.s. |
| Cycles of resubmission | 1 | 1-2 | 323 | 1 | 1-2 | 325 | n.s. |
| Days from resubmission to final decision | 31 | 15.5-58 | 323 | 34 | 16-59 | 325 | n.s. |

#### Compliance by Country

There were no differences in compliance between control and intervention groups across any corresponding author country of origin. Although we did not set out to compare differences in compliance with different ARRIVE subitems across countries, we present these data in Figure 3.

**Figure 3:**
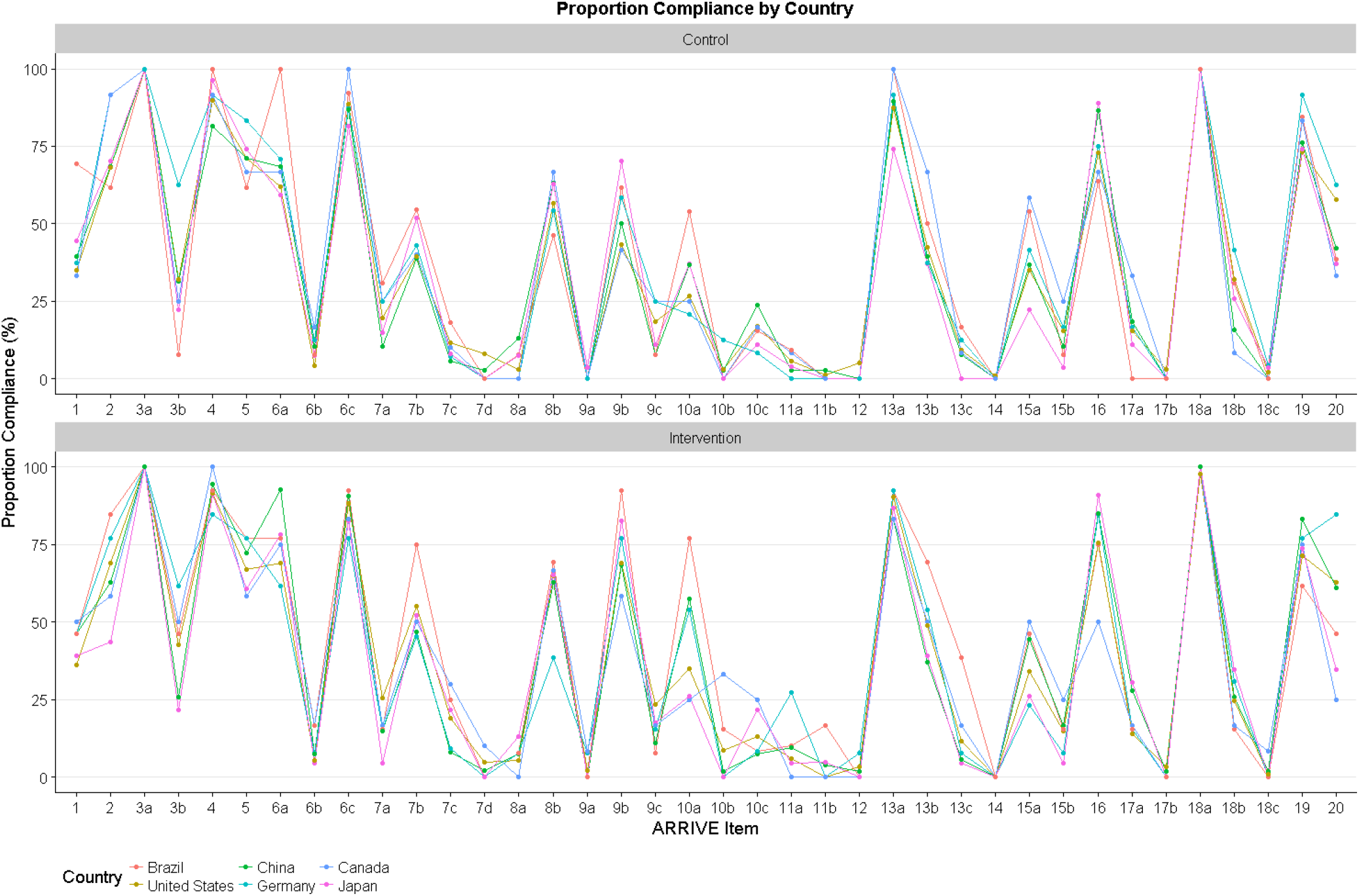
Compliance by country; Percentage compliance for each ARRIVE Item for manuscripts in each country (for countries with N manuscripts ≥10)

#### Human Studies Compliance

In manuscripts without human subjects, reporting of one ARRIVE subitem, 9b (husbandry related) increased significantly from 52.4% (172/316) in the control group to 76.9% (227/295) in the intervention group (X^2^ = 33.2, df=1, p<0.0001). In manuscripts containing human subjects, compliance also rose from 20.8% (5/24) to 51.35% (19/37) in the intervention group for this subitem, although we were limited by small sample sizes and this change was not found to be significant (X^2^= 4.47, df=1, p=1).

### Exploratory Outcomes

#### Compliance in True Intervention Group

Despite allocation to the intervention group, a small subset (n=31/332) of authors did not comply with the request to submit a completed checklist and therefore 31 manuscripts were in the intervention group without a completed ARRIVE checklist. We sought to determine compliance with each of the 38 subitems in the “true” intervention group (those submitted with a completed checklist), compared to the control group. The pattern of compliance is similar to that of the full intervention group compared to controls, suggesting that these instances of non-compliance did not impact on results. Summary statistics are presented in Table 3.

#### Landis Item Individual Compliance

In the ARRIVE guidelines, randomisation and blinding are part of the same subitem therefore a paper should report both randomisation and blinding to be compliant. However, to determine if there had been any changes in individual Landis items we investigated randomisation, blinding, and the other two Landis criteria items (reporting of a sample size calculation, reporting of exclusions) separately (Figure 4). Although we did not analyse these comparisons statistically, there appears to be some improvements in reporting of randomisation and sample size calculations. 29.1% (91/313) of manuscripts in the control group reported whether or not random assignment occurred, compared to 41.5% (125/301) in the intervention group (Cohen’s H effect size = 0.26). While 3.5% (12/40) of control manuscripts reported sample size calculations compared to 7.6% (25/330) in the intervention group (Cohen’s H effect size = 0.18). For the reporting of animal exclusions, 12.6% (43/340) of manuscripts complied in the control group versus 14.5% (48/332) in the intervention group. Finally, 18.8% (63/334) and 19.2% (62/323) of manuscripts reported blinded outcome assessment in the control and intervention groups respectively.

**Figure 4:**
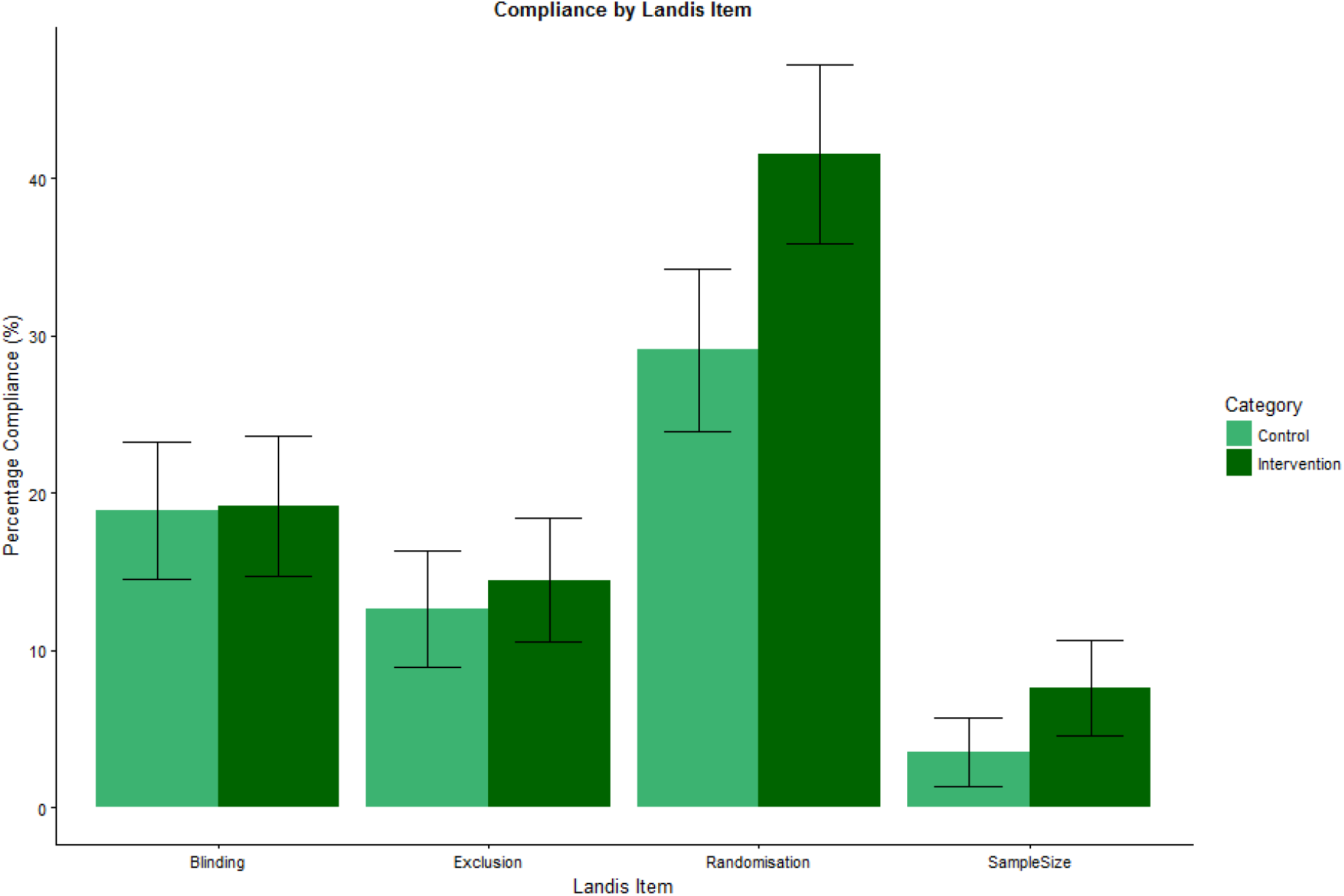
Landis 4 individual compliance; Percentage compliance for each Landis criteria item with 95% confidence intervals;

## Discussion

Requesting completion of an ARRIVE checklist at submission did not increase full adherence with the ARRIVE guidelines. Compliance with the operationalised ARRIVE checklist was poor overall, with no papers in either group even approaching full compliance; the median compliance was less than 40%, equivalent to around 15 of 38 items; and the intervention only increased compliance with one item, reporting of animal husbandry conditions. There is considerable room for improvement, and this study shows that an editorial policy of making ARRIVE checklist completion “mandatory” without compliance checks has little or no impact.

It may be that simply requesting that authors complete checklist, without any additional editorial checks to determine whether the checklist is truly indicative of compliance, may not be enough to improve adherence to the ARRIVE guidelines. Adherence to the reporting guidelines within in the clinical literature such as CONSORT and STROBE have been widely assessed and may inform interventions to improve compliance with preclinical guidelines. Journal endorsement of these guidelines appear to have improved reporting quality (Prady et al., 2008, Turner et al., 2012), however, it is often unclear what actions journals take to promote adherence (Stevens et al., 2014) and the extent of editorial involvement is likely to have an impact. Prior reports indicate that assessing compliance with reporting guidelines at the stage of peer review leads to a significant improvement of reporting quality (Cobo et al., 2011). Other approaches (e.g. actions on the part of funders or institutions) may also be beneficial, but a successful strategy is likely to be multi-dimensional. Further, the findings reported here and the limited agreement between outcome assessors both in this study and in the recent investigation of study quality following the introduction of a new editorial policy at *Nature* journals (Macleod, 2017) suggests that an important part of guideline development should be refinement of the content; the number of items (with fewer generally being better) and the agreement between assessors.

Our findings are in line with prior reports that endorsement by editors and reviewers has not significantly improved reporting of ARRIVE quality items (Baker et al., 2014, Gulin et al., 2015). We need therefore a better understanding of the barriers to implementing quality checklists for animal experiments. It has been suggested that requesting checklist adherence at the submission stage may be too late, given the observed correlation between reporting at the planning application stage and at the publication stage (Vogt et al., 2016). The PREPARE (Planning Research and Experimental Procedures on Animals: Recommendations for Excellence) guidelines (Smith et al., 2018) were published recently and may be a useful tool, in combination with the ARRIVE checklist, to promote a greater focus on experimental rigour at all stages of the research cycle.

Our results contrast with recent reports of improvement in quality following mandated checklist completion following a change in editorial policy at *Nature* journals (Han et al., 2017, Macleod, 2017). However, in both reports study quality was retrospectively assessed in publications published prior to and after the introduction of the Nature quality checklist, which was established in 2015 as part of an organisation wide approach with substantial editorial involvement. In contrast, the current trial investigated an intervention targeted at selected manuscripts, without further editorial involvement. Perhaps unsurprisingly, due to the additional time required for ARRIVE checklist requests, both the number of days manuscripts spent in the PLOS editorial office and the number of days from manuscript submission to AE assignment were found to be significantly longer in the intervention group. The editorial resource required to ensure that all accepted publications meet the requirements of the ARRIVE checklist is likely to be considerable, given that PLOS ONE is a high-volume publisher, with around 44,000 submissions per year. The most feasible and effective way to encourage compliance to the ARRIVE guidelines, or indeed any reporting guideline, remains to be determined, but an ongoing review of interventions to improve adherence to reporting guidelines may shed some light on this issue and direct future investigations (Blanco et al., 2017).

Another consideration is the perceived clarity of the checklist to authors and reviewers. Although reviewer agreement was generally high, a few questions were less well understood by our outcome assessors which suggests the current guidelines may require clearer dissemination among the research community.

### Limitations

Due to modest sample sizes we were unable to investigate whether the intervention was more successful in countries with high awareness and adoption of the ARRIVE guidelines such as the United Kingdom, where the ARRIVE guidelines were developed and where many institutions have endorsed them. Furthermore, we did not perform a power calculation for outcomes beyond our primary and main secondary outcomes and it is possible that this, coupled with stringent adjustments for multiplicity of testing in some instances, may have prevented us from detecting any significant differences.

Furthermore, our intervention only involved requests for authors to complete an ARRIVE checklist. PLOS ONE did not fully mandate checklist completion, as manuscripts without a checklist were still allowed to proceed through the trial. Furthermore, PLOS ONE did not evaluate the accuracy of the completed checklists against each manuscript. It is possible that further emphasis on evaluation and checklist adherence may result in an enhancement of study quality.

Our interpretation of compliance was also influenced by our operationalisation of the ARRIVE checklist used for outcome assessment. It was often difficult to determine how many of the details provided in the ARRIVE guidelines were sufficient for full compliance to that ARRIVE item.

There were unforeseen difficulties in attaining data for some outcomes, which meant that we could not assess all outcomes presented in our study protocol. This was most apparent for feasibility outcomes, where there were substantial deviations from our protocol. Furthermore, the project was subject to research waste due to overpowering our primary and secondary outcome measures. As manuscripts submitted to PLOS ONE as part of the study were treated differently, the existence of the study could have leaked to external sources however, to the best of our knowledge, this was not the case.

Since the ARRIVE guidelines were developed by the NC3Rs, no individual employed by the NC3Rs was permitted to conduct any outcome assessment on the IICARus platform.

### Conclusions

Research must be described in sufficient detail to allow research users critically to appraise experimental design, to allow them to assess the validity of the findings presented. Replication studies require, for their design, full details of what was done. Transparency in the reporting of research is paramount. Manuscripts must therefore be described in enough detail for readers to understand the research methodology and make informed judgement of quality and risk of bias. At present, reporting quality is, on average, disappointingly poor. However, our findings show that simply requesting that researchers improve reporting is not effective. It may be that a more formal adoption of research improvement strategies, with an original focus on a smaller number of items judged by a stakeholder to be of greatest importance, will allow an incremental approach to enabling and measuring improvement. Furthermore, editorial checks of compliance and further measures to mandate checklist completion may be required to see improvements in quality.

---

**Appendix 1:** Operationalised ARRIVE checklist for IICARus platform; Question Number, operationalised checklist number; questions not referred to explicitly in the ARRIVE guidelines are shown in grey.

| Section Title | Question Number | IICARus Question | ARRIVE Guidelines |
| --- | --- | --- | --- |
| **TITLE / ABSTRACT** |  |  |  |
| Title |  |  | **ARRIVE Item 1** |
|  | **1.1** | Is the species of animal model studied reported in the title? | Provide as accurate and concise a description of the content of the article as possible. |
|  | **1.2** | Is the biological mechanism, disease or pathophysiology studied, reported in the title? |  |
|  | **1.3** | Is the intervention or exposure reported in the title? |  |
| Abstract |  |  | **ARRIVE Item 2** |
|  | **2.1** | Is the objective or hypothesis reported in the abstract? | Provide an accurate summary of the background, research objectives, including details of the species or strain of animal used, key methods, principal findings and conclusions of the study. |
|  | **2.2** | Is the biological mechanism, disease or pathophysiology studied, reported in the abstract? |  |
|  | **2.3** | Is the intervention or exposure reported in the abstract? |  |
|  | **2.4** | Is the species or strain studied stated anywhere in the abstract? |  |
|  | **2.5** | Are the key methods of the study briefly summarised? |  |
|  | **2.6** | Are the principal findings of the study briefly summarised? |  |
|  | **2.7** | Are the conclusions of the study briefly summarised? |  |
|  | **0.1** | What animal species are used in this research? | Not mentioned in ARRIVE guidelines |
|  | **0.2** | Does the manuscript include human study? | Not mentioned in ARRIVE guidelines |
| **INTRODUCTION** |  |  |  |
| Background |  |  | **ARRIVE Item 3a** |
|  | **3.1** | Do the authors refer to previous work in the literature relating to this field? | Include sufficient scientific background (including relevant references to previous work) to understand the motivation and context for the study, and explain the experimental approach and rationale. |
|  |  |  | **Arrive Item 3b** |
|  | **3.2** | Is a statement reported about the rationale for using that animal species or animal disease model to address the scientific objectives? | Explain how and why the animal species and model being used can address the scientific objectives and, where appropriate, the study's relevance to human biology |
|  | **3.3** | If applicable to the research question, is there a statement describing the relevance of the study to human biology? |  |
| **Objectives** |  |  | **ARRIVE Item 4** |
|  | **4.1** | Is the objective or hypothesis reported in the introduction? | Clearly describe the primary and any secondary objectives of the study, or specific hypotheses being tested. |
| **METHODS** |  |  |  |
| **Ethical statement** |  |  | **ARRIVE Item 5** |
|  | **5.1** | Does the manuscript include an explicit statement of approval? | Indicate the nature of the ethical review permissions, relevant licences (e.g. Animal [Scientific Procedures] Act 1986), and national or institutional guidelines for the care and use of animals, that cover the research. |
|  | **5.2** | Does the manuscript identify the committee(s) approving the study protocol? |  |
|  | **5.3** | Does the manuscript name the international, national or institutional guidelines followed? |  |
|  | **5.4** | Does the manuscript report a protocol / permit number? |  |
| **Study Design** |  |  | **ARRIVE Item 6a** |
|  | **6.1** | Are the total number of experimental and control groups reported? | For each experiment, give brief details of the study design including: The number of experimental and control groups. |
|  |  |  | **ARRIVE Item 6c** |
|  | **6.2** | Is the experimental unit stated? |  |
|  | **6.3** | If the experimental unit is not stated is it clear what it is? | For each experiment, give brief details of the study design including: The experimental unit (e.g. a single animal, group or cage of animals). |
|  |  |  | **ARRIVE Item 6b** |
|  | **6.4** | Is randomisation reported? | For each experiment, give brief details of the study design including: Any steps taken to minimise the effects of subjective bias when allocating animals to treatment (e.g. randomisation procedure) and when assessing results (e.g. if done, describe who was blinded and when). |
|  | **6.5** | Does the manuscript include a statement about randomisation even if no randomisation was done? |  |
|  | **6.6** | Are assessors blinded for at least one of the outcomes measured? |  |
|  | **6.7** | Does the manuscript include a statement about blinding even if no blinding was done? |  |
| Experimental Procedures |  |  | **ARRIVE Item 7a** |
|  | **7.1.1** | Vehicle(s) reported? | For each experiment and each experimental group, including controls, provide precise details of all procedures carried out. For example: a. How (e.g. drug formulation and dose, site and route of administration, anaesthesia and analgesia used [including monitoring], surgical procedure, method of euthanasia). Provide details of any specialist equipment used, including supplier(s). |
|  | **7.1.2** | Vehicle volume(s) reported? |  |
|  | **7.1.3** | Intervention/exposure dose(s) reported? |  |
|  | **7.1.4** | Route(s) of administration reported? |  |
|  | **7.1.5** | Site(s) of administration reported? |  |
|  | **7.1.6** | Frequency of administration reported? |  |
|  | **7.1.7** | Supplier(s) reported? |  |
| Experimental Procedures (Control) |  |  |  |
|  | **7.3.1** | Is the control reported? |  |
|  | **7.3.2** | Is the control dose or volume reported? |  |
|  | **7.3.3** | Is the control route reported? |  |
|  | **7.3.4** | Is the control site of administration reported? |  |
|  | **7.3.5** | Is the frequency of administration reported? |  |
|  | **7.3.6** | If a control (e.g. sham) surgical procedure was carried out do they describe the methods used? |  |
| **Surgery and Anaesthesia** |  |  |  |
|  | **7.5.1** | Is surgical anaesthesia use reported? |  |
|  | **7.5.2** | Is the anaesthesia route reported? |  |
|  | **7.5.3** | Is the anaesthetic reported? |  |
|  | **7.5.4** | Is the anaesthesia dose reported? |  |
|  | **7.5.5** | Are the methods used for surgical procedures clearly described? |  |
|  | **7.5.6** | Are the suppliers for any specialist surgical equipment reported? |  |
|  | **7.5.7** | Is the monitoring of at least one physiological parameters during surgical anaesthesia reported? |  |
|  | **7.5.8** | Is the use of an analgesic, or a reason why analgesic was not used, reported? |  |
| **Euthanasia** |  |  |  |
|  | **7.6.1** | Is euthanasia, sacrifice etc. reported? |  |
|  | **7.6.2** | Is the method of euthanasia reported? |  |
|  | **7.1.8** | If a surgical procedure was carried out was it part of model induction? | Not mentioned in ARRIVE guidelines |
|  | **7.1.9** | If a surgical procedure was carried out was it part of either treatment or outcome measurement(s)? | Not mentioned in ARRIVE guidelines |
| **Experimental Procedures** |  |  | **ARRIVE Item 7b** |
|  | **7.2.1** | Does the manuscript describe when the intervention/exposure group procedures were carried out? | For each experiment and each experimental group, including controls, provide precise details of all procedures carried out. For example: When (e.g. time of day). |
| Experimental Procedures (Control) |  |  |  |
|  | **7.4.1** | Does the manuscript describe when the control/comparator intervention procedures were carried out? |  |
| Experimental Procedures |  |  | **ARRIVE Item 7c** |
|  | **7.2.2** | Does the manuscript describe where the intervention/exposure group procedures were carried out? | For each experiment and each experimental group, including controls, provide precise details of all procedures carried out. For example: Where (e.g. home cage, laboratory, water maze). |
| Experimental Procedures (Control) |  |  |  |
|  | **7.4.2** | Does the manuscript describe where the control/comparator intervention procedures were carried out? |  |
| Experimental Procedures |  |  | **ARRIVE Item 7d** |
|  | **7.2.3** | Is any rationale for the use of the intervention/exposure reported? | For each experiment and each experimental group, including controls, provide precise details of all procedures carried out. For example: Why (e.g. rationale for choice of specific anaesthetic, route of administration, drug dose used). |
| Experimental Procedures (Control) |  |  |  |
|  | **7.4.3** | Is any rationale for the use of the control/comparator group reported? |  |
| Experimental animals |  |  | **ARRIVE Item 8a** |
|  | 8.1 | Is the animal species reported? | Provide details of the animals used, including species, strain, sex, developmental stage (e.g. mean or median age plus age range) and weight (e.g. mean or median weight plus weight range). |
|  | 8.2 | Is the strain of the animals reported? |  |
|  | 8.3 | Is the sex of the animals reported? |  |
|  | 8.4 | Is the age of the animals reported? |  |
|  | 8.5 | Is the weight of the animals reported? |  |
|  |  |  | **ARRIVE Item 8b** |
|  | 8.6 | For studies using transgenic animals, do the authors report: 1) The genetic modification status (knockout, | Provide further relevant information such as the source of animals, |
|  |  | overexpression etc.), 2) The genotype (homozygous, heterozygous) and 3) The manipulated gene/s? | international strain nomenclature, genetic modification status (e.g. knock-out or transgenic), genotype, health/immune status, drug or test naïve, previous procedures, etc. |
|  | 8.8 | Is the source/supplier of the animals reported? |  |
| **Housing** |  |  | **ARRIVE Item 9a** |
|  | 9.1.1 | Is the biosecurity level of the facility reported? | Housing (type of facility e.g. specific pathogen free [SPF]; type of cage or housing; bedding material; number of cage companions; tank shape and material etc. for fish). |
|  | 9.1.2 | Is the type of cage or housing reported? |  |
|  | 9.1.3 | Is the bedding material reported? |  |
|  | 9.1.4 | Is the number of cage companions reported? |  |
|  | 9.2.4 | For experiments involving fish, are the tank dimensions or materials reported? |  |
| **Husbandry** |  |  | **ARRIVE Item 9b** |
|  | 9.2.1 | Are the light/dark cycle conditions reported? | Husbandry conditions (e.g. breeding programme, light/dark cycle, temperature, quality of water etc for fish, type of food, access to food and water, environmental enrichment).<br>for fish, type of food, access to food and water, environmental enrichment). |
|  | 9.2.2 | Is the temperature reported? |  |
|  | 9.2.3 | For experiments involving fish, is the quality of the water reported? |  |
|  | 9.2.5 | Is the type of food provided reported? |  |
|  | 9.2.6 | Are the conditions around access to food reported? |  |
|  | 9.2.7 | Are the conditions around access to drinking water reported? |  |
|  | 9.2.8 | Is any environmental enrichment reported? |  |
| **Welfare** |  |  | **ARRIVE Item 9c** |
|  | 9.3.1 | Have they reported any welfare assessment or intervention before, during, or after the experiment? | Welfare-related assessments and interventions that were carried out prior to, during, or after the experiment. |
| **Sample size** |  |  | **ARRIVE Item 10a** |
|  | 10.1 | Is the total number of animal used for the experiment reported? | Specify the total number of animals used in each |
|  | 10.2 | Is the number of animals in each experimental group reported? |  |
|  |  |  | ARRIVE Item 10b |
|  | 10.3 | Is a sample size calculation reported? | Explain how the number of animals was arrived at.<br>Provide details of any sample size calculation used. |
|  | 10.4 | Is the statistical method for the sample size calculation reported or any other explanation provided? |  |
|  |  |  | ARRIVE Item 10c |
|  | 10.5 | Is the number of independently replicated experiments reported? | Indicate the number of independent replications of each experiment, if relevant |
|  | 11.1 | Is allocation concealment reported? | Not mentioned in ARRIVE guidelines |
| Allocating animals to experimental groups |  |  | ARRIVE Item 11a |
|  | 11.2 | Are the methods of allocation to group (i.e. randomisation, matching) described? | Give full details of how animals were allocated to experimental groups, including randomisation or matching if done. |
|  |  |  | ARRIVE Item 11b |
|  | 11.3 | Is the order in which animals receive treatments defined? | Describe the order in which the animals in the different experimental groups were treated and assessed. |
|  | 11.4 | Is the order in which outcomes are assessed in different animals reported? |  |
| Experimental outcomes |  |  | ARRIVE Item 12 |
|  | 12.1 | Are outcomes reported identified as being either primary or secondary? | Clearly define the primary and secondary experimental outcomes assessed (e.g. cell death, molecular markers, behavioural changes). |
|  | 12.2 | Is at least one outcome measure described? |  |
| Statistical methods |  |  | ARRIVE Item 13a |
|  | 13.1 | Is at least one outcome measure associated with at least one statistical test? | Provide details of the statistical methods used for each analysis. |
|  |  |  | ARRIVE Item 13b |
|  | 13.2 | Is the unit of analysis for at least one tests explicitly specified? | Specify the unit of analysis for each dataset (e.g. single animal, group of animals, single neuron). |
|  |  |  | ARRIVE Item 13c |
|  | 13.3 | Does the publication include a method to assess whether the data meet the assumptions of the statistical tests used? | Describe any methods used to assess whether the data met the assumptions of the statistical approach. |
| RESULTS |  |  |  |
| Numbers analysed |  |  | ARRIVE Item 14 |
|  | 8.7 | Are the animals used in the study reported to be drug or test naïve prior to treatment or testing? | For each experimental group, report relevant characteristics and health status of animals (e.g. weight, microbiological status, and drug or test naïve) prior to treatment or testing. (This information can often be tabulated). |
|  | 8.9 | Is the health status of the animals reported? |  |
|  |  |  | ARRIVE Item 15a |
|  | 14.1 | Is the number of animals for each group reported for each analysis? | Report the number of animals in each group included in each analysis. Report absolute numbers (e.g. 10/20, not 50%2). |
|  |  |  | ARRIVE Item 15b |
|  | 14.2 | Are reasons for the exclusion of animals (for any outcome) given? | If any animals or data were not included in the analysis, explain why. |
|  |  |  | ARRIVE Item 16 |
|  | 15.1 | Are findings presented with a measure of precision? | Report the results for each analysis carried out, with a measure of precision (e.g. standard error or confidence interval). |
|  | 15.2 | Is the measure of precision defined? |  |
|  |  |  | ARRIVE Item 17a |
|  | 16.1 | Is there a statement indicating whether or not adverse events occurred for at least one experimental group? | Give details of all important adverse events in each experimental group. |
|  |  |  | ARRIVE Item 17b |
|  | 16.2 | Are any modifications to the experimental design to reduced adverse effects reported? | Describe any modifications to the experimental protocols made to reduce adverse events. |
| **DISCUSSION** |  |  |  |
| Interpretation/<br>scientific<br>implications |  |  | ARRIVE Item 18a |
|  | 17.1 | Are the results interpreted in the context of the study hypothesis or objectives? | Interpret the results, taking into account the study objectives and hypotheses, current theory and other relevant studies in the literature. |
|  | 17.2 | Are the results interpreted in the context of other studies in the literature? |  |
|  |  |  | ARRIVE Item 18b |
|  | 17.3 | Are the limitations of the study design and/or execution discussed? | Comment on the study limitations including any potential sources of bias, any limitations of the animal model, and the imprecision associated with the results. |
|  |  |  | ARRIVE Item 18c |
|  | 17.4 | Are any implications of the experimental methods or findings for the replacement, refinement or reduction (the 3Rs) of the use of animals in research discussed? | Describe any implications of your experimental methods or findings for the replacement, refinement or reduction (the 3Rs) of the use of animals in research. |
| Generalisability/<br>translation |  |  | ARRIVE Item 19 |
|  | 18.1 | Is there a statement about how the findings of this study might translate to other species or systems, such as any relevance to human biology? | Comment on whether, and how, the findings of this study are likely to translate to other species or systems, including any relevance to human biology. |
| Funding |  |  | ARRIVE Item 20 |
|  | 19.1 | Do the authors report funding source(s)? | List all funding sources (including grant number) and the role of the funder(s) in the study. |
|  | 19.2 | Do the authors include the grant number (grant #)? |  |
|  | 19.3 | Has the role of the funders been reported? |  |
|  | 19.4 | Is there a statement of competing/conflict of interests? | Not mentioned in ARRIVE guidelines |

